# Habitat choice stabilizes metapopulation dynamics through increased ecological specialisation

**DOI:** 10.1101/267575

**Authors:** Frederik Mortier, Staffan Jacob, Martijn L. Vandegehuchte, Dries Bonte

## Abstract

Dispersal is a key trait responsible for the spread of individuals and genes among local populations, thereby generating eco-evolutionary interactions. Especially in heterogeneous metapopulations, a tight coupling between dispersal, population dynamics and the evolution of local adaptation is expected. In this respect, current theory predicts dispersal to counteract ecological specialisation by redistributing locally selected phenotypes (i.e. migration load). However, in nature we observe that some specialists exhibit a strong dispersal capacity.

Habitat choice following informed dispersal decisions, provides a possible mechanism for individuals to match the environment to their phenotype, thereby enabling the persistence of evolved ecological specialisation. How such informed decisions affect the evolution of dispersal and ecological specialisation and how these, in turn, influence metapopulation dynamics is yet to be determined.

By means of individual-based modelling, we show that informed decisions on both departure and settlement decouples the evolution of dispersal and generalism, favouring highly dispersive specialists. Choice at settlement decouples dispersal from ecological specialisation most effectively. Additionally, habitat choice stabilizes local and metapopulation demography because of the maintenance of ecological specialisation at all levels of dispersal propensity.

We advocate considering habitat choice in spatially structured ecological models to improve demographic predictions in the face of environmental change.

## Introduction

Most populations are spatially structured and organized in metapopulations. Local populations are connected by dispersal, the movement of individuals or propagules that potentially generates gene flow across space [1]. Dispersal thus acts as a glue that links local gene pools, local population dynamics and metapopulation dynamics [2]. Dispersal is known to evolve in response to metapopulation structure as a bet-hedging strategy in spatio-temporally variable environments [1,3–5]. It is also known to evolve in response to local drivers, for instance to escape kin competition and inbreeding depression [1, 3]. Dispersal is thus an essential attribute for fitness maximisation [6]. Ultimately, these benefits are balanced against dispersal costs to determine the optimal dispersal strategy-in terms of frequency and distance.

In metapopulations, habitat heterogeneity introduces additional costs to dispersal. In such a setting, divergent selective pressures among local habitats can result in local adaptation [7]. Often, local adaptation comes at the cost of lower performance in other environments, which can lead populations to adapt to a smaller subset of available habitats-i.e. ecological specialisation [8]. If specialists experience the landscape as highly heterogeneous when dispersing [9], they are more likely to end up in unsuitable habitat. Ecological specialisation is therefore predicted to select against dispersal [10, 11]. Reciprocally, dispersal hinders local adaptation by mixing gene pools and thus opposes the evolution of specialisation [12–15]. This trade-off between dispersal and local adaptation is expected to maximize performance across an environmentally heterogeneous landscape, and thus homogenize fitness. High dispersal rates and ecological specialisation are, therefore, difficult to reconcile (theoretical evidence: [11,15–19], empirical evidence: [12,13,20]).

If dispersal involves habitat choice, dispersal and ecological specialisation may be reconcilable. From a pure movement perspective, optimal foraging theory inherently implements choice-based movement, with individuals weighing the cost of foraging time against the benefit of finding the optimal resource [21]. Such adaptive behaviour could also be relevant in the context of dispersal-when considering movements directly linked to reproduction. Rather than assuming that each member of a population disperses randomly with the same probability, informed dispersal implies a non-random subset of the local population dispersing and/or dispersers being redistributed in a non-random way across the landscape [22–25]. Individuals can indeed use information about their phenotype and environment to decide whether to disperse (departure decision), and where to go (settlement decision; [26]). Several organisms gather and use information during movement [27–29]. Intuitive in the light of evolution, habitat choice enables individuals to track the environment that best matches their phenotype while also reaping the benefits of dispersal [30, 31]. Furthermore, theoretical and experimental studies show how habitat choice cascades into further life-history evolution, increasing local adaptation and ecological specialisation (Theory: [32–36]; empirical: [22, 37]).

With habitat choice, individuals integrate information to preferentially disperse towards habitat that maximizes their fitness, which affects the evolution of dispersal and specialisation. Subsequently, rapid evolutionary changes are expected to affect ecological dynamics in the same time frame according to the eco-evolutionary framework [38]. Understanding changes in metapopulation dynamics should enable us to understand and predict metapopulation persistence in a spatially structured and heterogeneous context. Here, we develop an individual-based model to study the impact of habitat choice on the evolution of dispersal and ecological specialisation. We separately analyse 1) how dispersal and habitat choice affect the evolution of ecological specialisation, and 2) how ecological specialisation and habitat choice affect dispersal evolution. Furthermore, we quantified the consequences of these evolutionary processes for metapopulation dynamics. While random dispersal should lead to a trade-off between dispersal and ecological specialisation, we predict habitat choice to favour specialists, even with high levels of dispersal. Additionally, we predict that habitat choice favours high levels of dispersal, even in specialists. Moreover, dispersal can affect how local populations within a metapopulation vary over time, increasing or decreasing metapopulation stability and synchrony [39–41]. We predict that the individual fitness maximisation by habitat choice should likewise favour the metapopulation by resulting in an increased metapopulation size and stability. Finally, since the consequences of habitat choice for ecological and evolutionary dynamics have been predicted to differ depending on the timing of the informed decision [42], we modelled four dispersal scenarios: random dispersal, habitat choice at departure, habitat choice at settlement, and habitat choice at both departure and settlement.

## Model

### Landscape

We model a finite landscape: a toroidal lattice of 32 × 32 grid cells. Each patch (i.e. grid cell) has a random environmental value υ_x,y_ ∈ [0, 1] at coordinates x,y, without any spatial autocorrelation. This environmental value is the local selective pressure. Its values are randomly distributed in space and constant in time creating a heterogeneous environment. Additionally, each patch contains a certain amount of resources (*G*_*x,y*_) that regulate local consumer population densities.

### Population

For simplicity, we model an asexually reproducing organism with discrete generations. The sequence of life-history events for each individual starts with dispersal and is succeeded by reproduction, then population regulation. This closely resembles soft selection in a semelparous species with a single dispersal phase [43]. These life-history events are explained below and assumed parameters are summarized in table 1.

#### Evolving traits

We model three traits that can be allowed to evolve:

The **optimal habitat trait (*muT*)** determines the optimum of the environmental value (υ_x,y_) for an individual to have its highest possible fitness (*muT* ∈ [0,1]).

The **niche width (*varT*)** determines the extent of ecological specialisation, by determining an individual’s fitness for values of the environment (υ_x_,_y_) a certain distance away from the individual’s optimal habitat (*muT*). A wide niche results in higher fitness far away from the optimal habitat (eq. 2; derived from Chaianunporn & Hoverstad [19], but decreases fitness in the optimal habitat (eq. 3, fig. 1). The match of an individual’s optimal habitat (*muT*) with the local environmental value (υ_x,y_) combined with its *varT* determines the individual’s efficiency in this particular habitat (α_i_, eq. 2). The amount of gathered resources (*F*_*i*_) combines the individual’s efficiency (α_*i*_) with the local resource density (*R*_*x,y*_) and is proportional to the individual’s expected reproductive success (eq. 1).

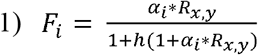

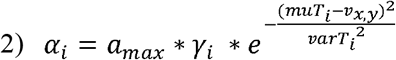

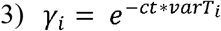

*F*_*i*_ represents the gathered resources by individual i, determined by a resource-consumer model with h being handling time, *R*_*xy*_ the amount of resources present locally, *a*_max_ the maximum encounter rate, *muT*_*i*_ and *varT*_*i*_ being the optimal habitat trait and niche width respectively for that individual and *υ*_*x,y*_ the local environmental value. γ_*i*_ implements the niche width-performance trade-off with *ct* indicating the strength of the trade-off.

**Figure 1:**
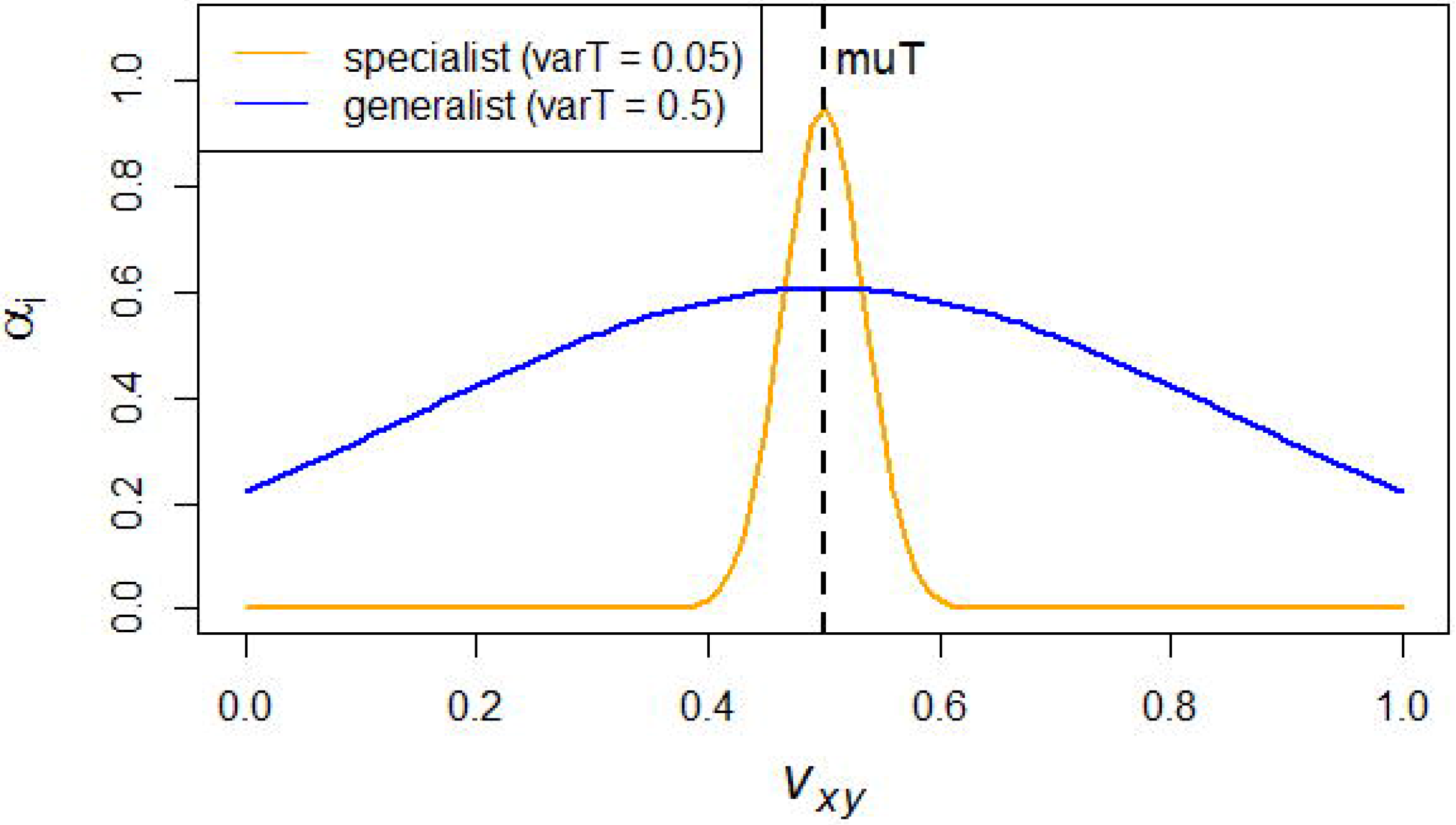
relation of local environmental value to individual efficiency in that location for a specialist (orange) and a generalist (blue) with optimal environment (muT) = 0.5.

The **dispersal trait (d)** represents the individual’s inclination to disperse. The role of this trait is explained in more detail in the next section.

#### Dispersal

Individuals disperse before selection occurs. We model two decision points in a dispersal event:

First, at **departure**, an individual disperses with a probability equal to its dispersal trait (d) if departure is random, meaning that a higher dispersal trait implies a higher tendency to disperse. With departure choice, the dispersal trait (d) represents the minimal acceptable local expected reproductive output at which an individual chooses not to disperse. Below this threshold, the local conditions are considered too bad and the individual leaves. In parallel with random departure, a higher threshold implies a higher tendency to disperse.

At **settlement**, an individual settles in a random patch within its dispersal range determined by a maximum dispersal distance (*dr*) if settlement is random (all patches 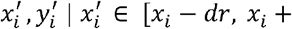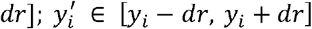. Its current location is excluded from this range to force dispersing individuals to change location. With habitat choice at settlement, the dispersing individual settles in the location where the local environmental value (*υ_x,y_*) best matches its own optimal habitat trait (*muT*) within its dispersal range.

Note that habitat choice at both decision points involves evaluating how well the individual’s optimal habitat (*muT*) trait matches the environmental value of a patch *υ_x,y_*) [24].

#### Reproduction

Reproducing individuals have an expected number of offspring (λ_*i*_) proportional to their gathered resources (*F*_*i*_) in the patch after the dispersal phase.

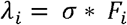

σ Indicates how many offspring each unit of resources results in. The actual reproductive output of an individual is sampled from a Poisson distribution with mean *λ_i_*.

#### Local population regulation

Local consumer populations are regulated through local resource availability (*G*_*xy*_). These resources are restocked each generation according to a logistic growth function,

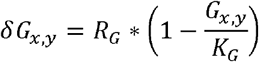

where the local resource increase *δG*_*x,y*_depends on *G*_*x,y*_, the amount of resources already present locally. Furthermore, *R*_*G*_ and *K*_*G*_ represent the optimal growth rate and carrying capacity of the resources respectively. Resources are consumed proportionally to the number of offspring. A consumer’s offspring without the required amount of resources in their local patch will die after depleting the leftover resources should there be any left. Population regulation depends on resource availability and, consequently, is density dependent, while habitat choice is decoupled from local population densities. Hereby, we avoid simulating density-dependent habitat choice (see [31] for a comparison of different modes of habitat choice).

#### Mutation

Non-fixed traits mutate at a rate of 0.01 generation^−1^. The optimal habitat trait (*muT*), dispersal trait (*d*) and niche width (*varT*) mutate by randomly sampling a new trait value from a normal distribution with the initial trait value as mean and standard deviation 0.1. New values of *muT* are limited to the range of possible environmental values [0, 1].

### Simulations

We analyse four models that represent all combinations of either random dispersal or habitat choice at departure and at settlement. First, we analyse how niche width (*varT*) evolves for different fixed values of the dispersal trait (d) and the different habitat choice scenarios. Second, we analyse how dispersal evolves for different fixed values of niche width with the different combinations of habitat choice. Fixed traits are varied over 20 values with equal increments within a range (i.e. random departure d:[0, 1], informed departure d: [0, 5], *varT:* [0, 0.5]). The range of the dispersal trait (d) in informed departure scenarios differs from the random scenarios since it represents the minimal acceptable reproductive output of an individual instead of its dispersal propensity. This range of d in the informed departure scenarios, however, results in actual dispersal propensities that cover the range [0, 1]. Each scenario is replicated 10 times with each replicate simulated for 500 generations. We analyse average niche width of the last generation in scenarios with evolving niche width, and the proportion of individuals that dispersed in the last generation in scenarios with the dispersal trait evolving. Additionally, we analyse the metapopulation size (i.e. total number of individuals) in the last generation and temporal variability in local population sizes, temporal variability in metapopulation size and asynchrony (α-, γ- and β-variability respectively [44],in the last five generations for each replicate.

#### Initialisation

We initialize each replicate by allocating 70000 individuals randomly across the 32 × 32-landscape grid. This initial metapopulation size close to the consumers’ carrying capacity avoids drift effects. Each individual’s optimal habitat trait value is sampled from a uniform distribution between 0 and 1.

Unless it was fixed for that scenario, niche width and the dispersal trait are randomly sampled from a uniform distribution (with the same range as their fixed values for the fixed scenarios).

#### Imperfect habitat choice

In addition to the analyses presented in this manuscript, we tested whether imperfect rather than perfect choice (at departure and settlement) leads to different eco-evolutionary dynamics (see *supplementary material 2*). We modelled imperfect choice as a probability, at each decision point, that the individual chooses randomly instead of in an informed way.

**Table 1:** Assumed parameters

|  |  |  |
| --- | --- | --- |
| $R_G$ | Optimal resource growth rate | 0.25 |
| $K_G$ | Resource carrying capacity | 1 |
| $h$ | Handling time | 0.2 |
| $a_{max}$ | Maximum encounter rate | 0.05 |
| $ct$ | Cost of generalism | 1 |
|  | mutation rate | 0.01 generation<sup>-1</sup> |
| $dr$ | Dispersal range | 2 |
| $\sigma$ | Resource conversion factor | 300 |

## Results

### Niche width evolution

With random dispersal (fig. 2: orange crosses), we find a gradual increase of niche width (i.e. ecological generalism) when increasing dispersal propensity. This means that, in accordance with classical predictions [11, 15], a low dispersal propensity favours specialism while a high dispersal propensity leads to the evolution of generalists. Habitat choice at departure results in a pattern comparable to that of the uninformed model, with a low level of dispersal in specialists or a high level of dispersal in generalists (fig. 2: orange circles). However, habitat choice at departure enables specialism to still evolve at a higher dispersal propensity than in scenarios of random dispersal. The inflexion point shifts from a dispersal propensity of around 0.35 to one of around 0.65. In contrast, habitat choice at settlement favours specialism in all scenarios (fig. 2: green crosses), indicating that specialism evolves regardless of dispersal propensity. Additionally, a departure decision results in a trend of even stronger specialism than if specialism evolves when a settlement decision is made. A combination of departure and settlement decision shows specialism evolving in all scenarios but more strongly so at lower dispersal propensities (fig. 2: green circles).

**Figure 2:**
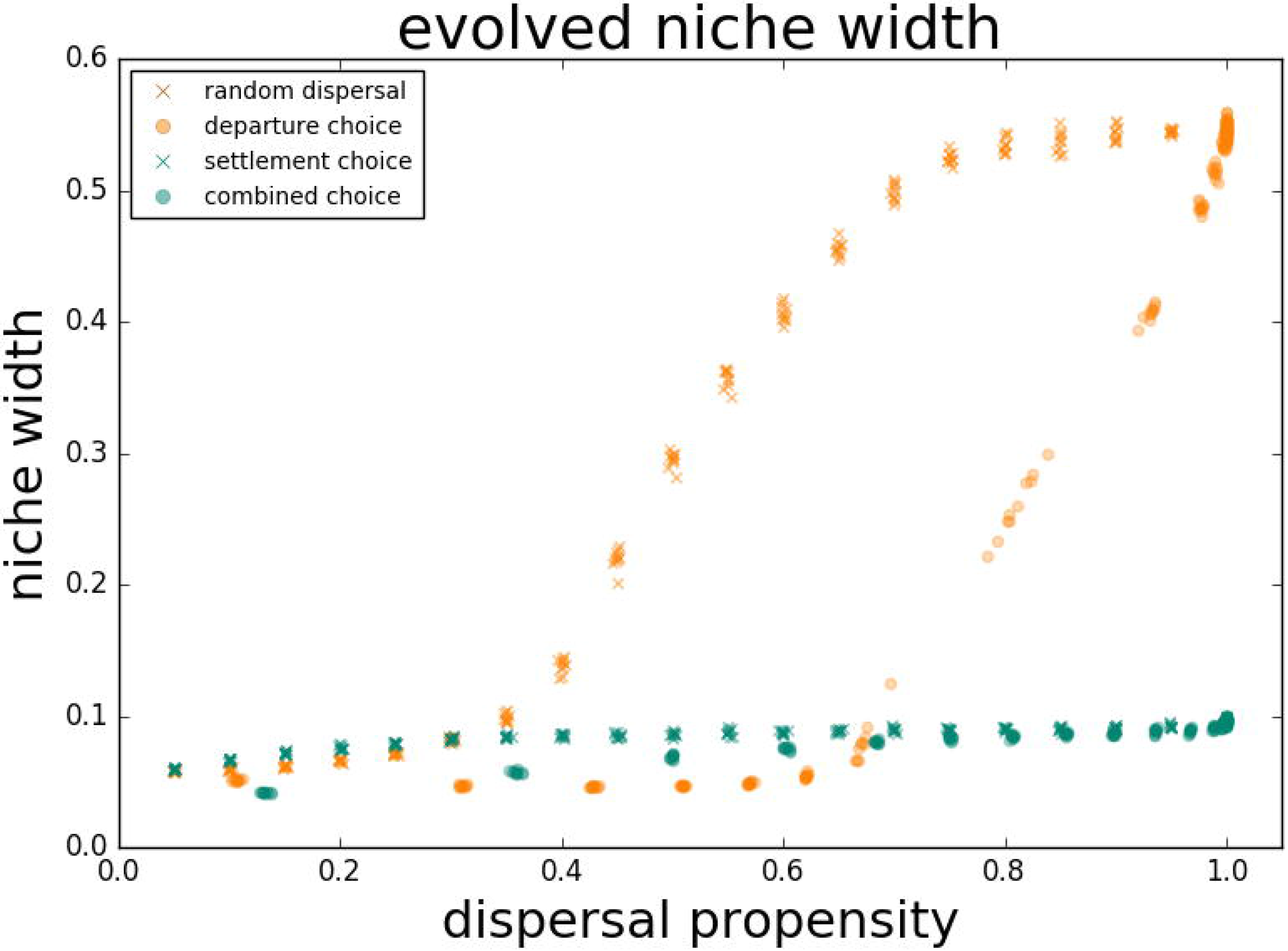
Niche width evolution in relation to effective dispersal propensity. Scenarios of random (×) and informed (○) departure combined with random (orange) or informed (green) settlement. Because the fixed dispersal trait does not equate the dispersal propensity in informed departure scenarios (○), their corresponding data points are not equally distributed across the dispersal propensity range.

Regarding ecological dynamics, we find that increasing dispersal propensity results in a decreased metapopulation size when dispersal is random or only informed at departure (fig. 3). Metapopulations exhibiting a low level of dispersal as well as those in which individuals have informed settlement achieve larger metapopulations (fig. 3) with more stable local populations (α, fig. 4). Note that those scenarios with lower metapopulation sizes correspond with scenarios in which generalism evolved. If dispersers only choose at departure, metapopulation sizes decrease to a minimum around the switch from specialism to generalism (fig. 3: orange circles). Also, the very dispersive scenarios with informed settlement result in local population variability being equal to or higher than without settlement decision, while metapopulation sizes are still markedly higher (fig. 3-4). Metapopulation variability (γ) and asynchrony (β) did not show clear trends (see *supplementary material 1*).

**Figure 3:**
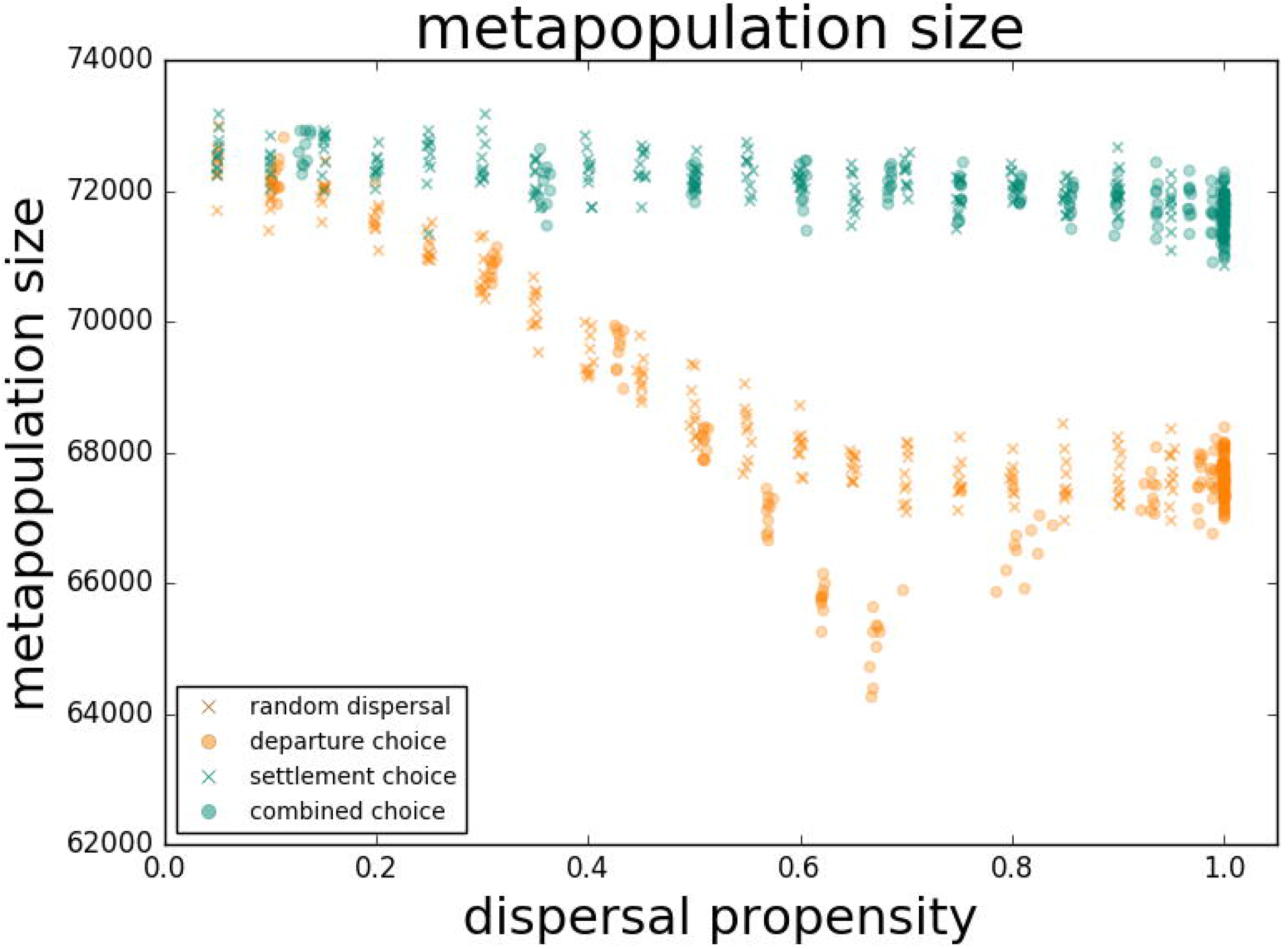
Metapopulation size for scenarios with fixed dispersal trait. Scenarios of random (×) and informed (○) departure combined with random (orange) or informed (green) settlement.

**Figure 4:**
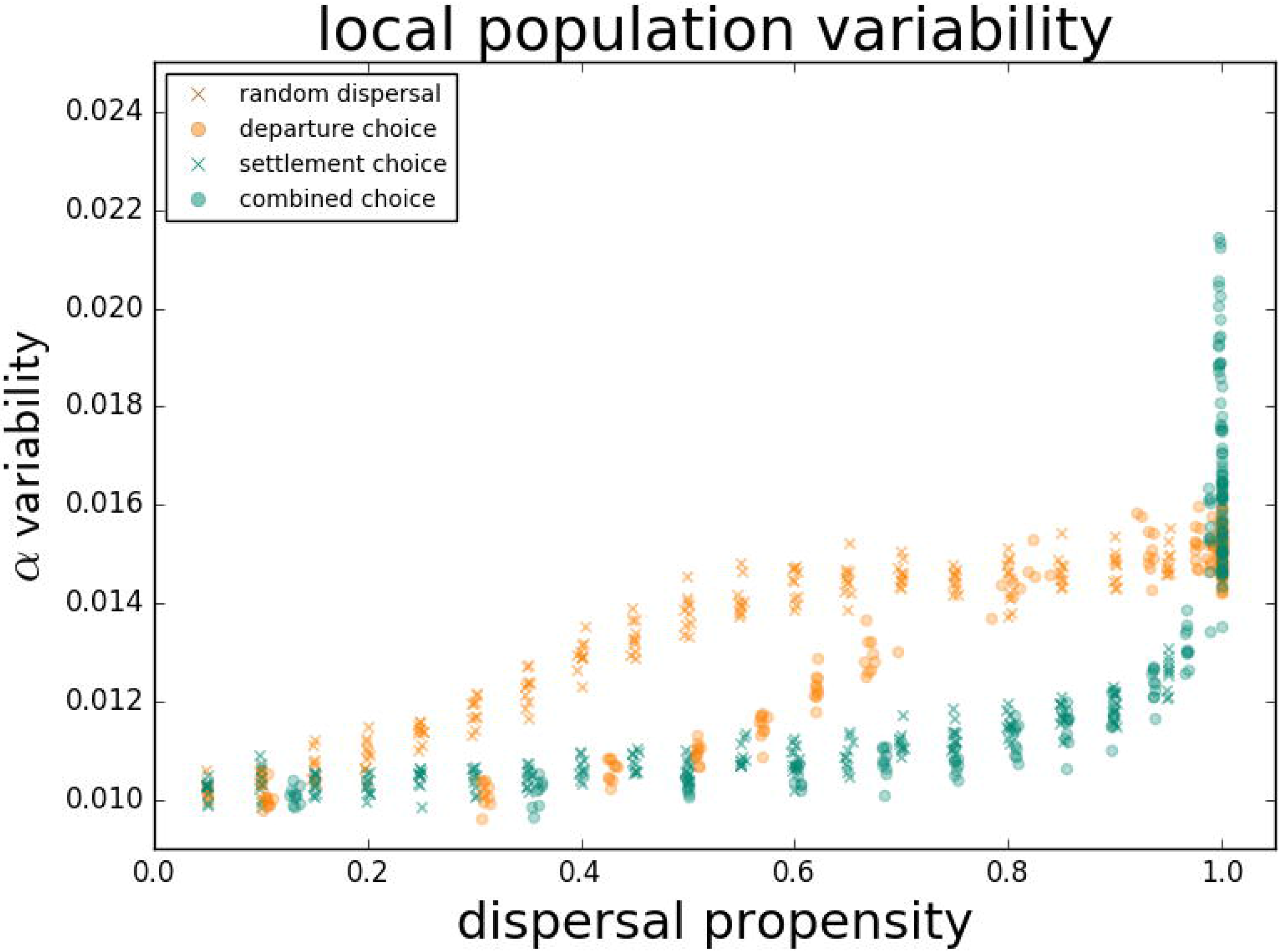
Local population variability for scenarios with fixed dispersal trait. Scenarios of random (×) and informed (○) departure combined with random (orange) or informed (green) settlement.

### Dispersal evolution

With random dispersal, dispersal propensity increases with niche width as predicted, but overall levels of dispersal propensity are low (fig. 5, left panel, orange crosses). With a departure decision, dispersal is also relatively infrequent (fig. 5, left panel, orange circles). In this case, however, dispersal propensity decreases with niche width, resulting in the most specialized individuals being the most dispersive (fig. 5, right panel, orange circles). With a departure choice, only mismatched individuals disperse (i.e. individuals whose optimal habitat trait differs significantly from the habitat value of their location). Since specialists only have a limited range of suitable habitats, they are less likely than generalists to find themselves in sufficiently suitable habitat. Therefore, specialists benefit more from dispersal with a departure decision. With a settlement decision, dispersal propensity is markedly higher (fig. 5, green crosses). Here, dispersal propensity drops slightly in metapopulations of the most specialized individuals (i.e. lowest niche width). With a decision both at departure and settlement, dispersal propensity is relatively high (fig. 5, green circles). It is slightly lower than in scenarios with only| a settlement decision because the additional departure decision prevents a proportion of the metapopulation from dispersing. In this scenario, only the very specialized individuals exhibit a significant decrease in tendency to disperse.

**Figure 5:**
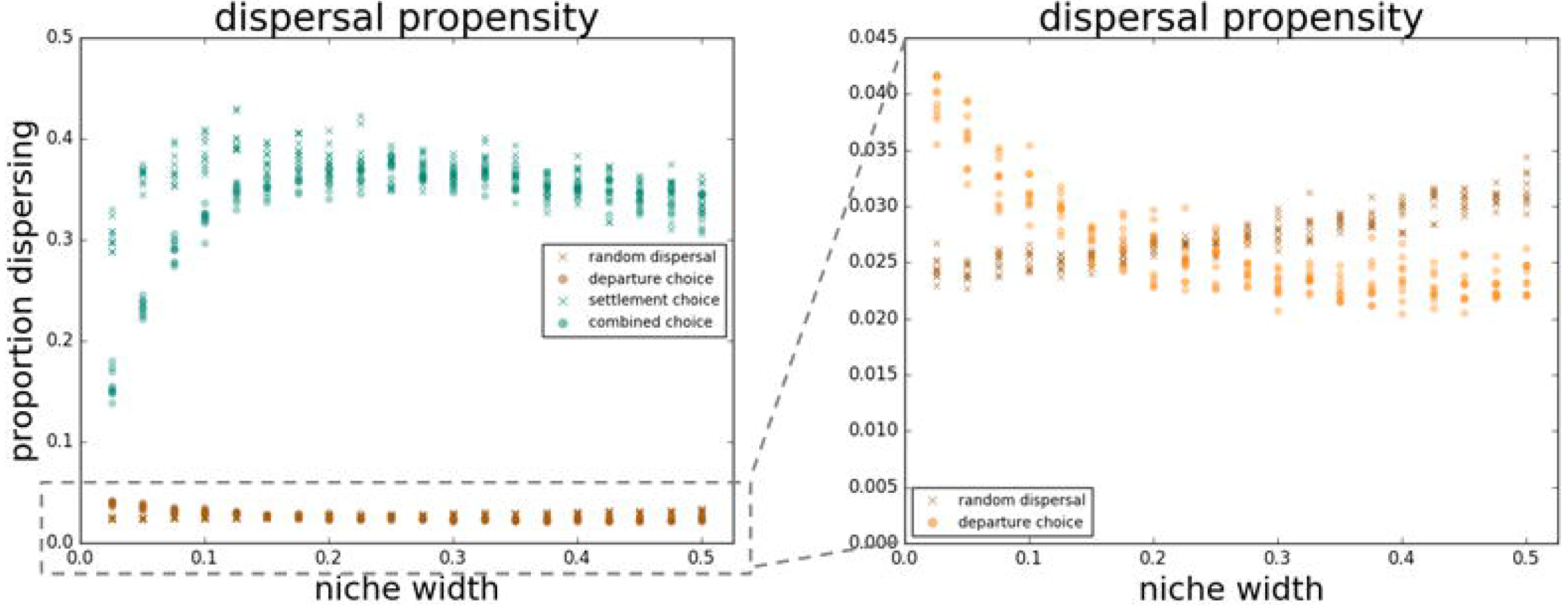
Effective dispersal propensity resulting from an evolving dispersal trait under fixed levels of niche width. Scenarios of random (×) and informed (○) departure combined with random (orange) or informed (green) settlement. Right panel zooms in on the scenarios of random settlement.

With increasing niche width, metapopulation size decreases (fig. 6) and local population variability increases (α, fig. 7). We find no noteworthy effect of the departure decision nor the settlement decision on metapopulation sizes. Local population variability does not show a strong effect of any of the habitat choice scenarios either, except for the slightly more variable local populations with a settlement choice. Metapopulation variability (γ) and asynchrony (β) did not show clear trends (see *supplementary material 1*).

**Figure 6:**
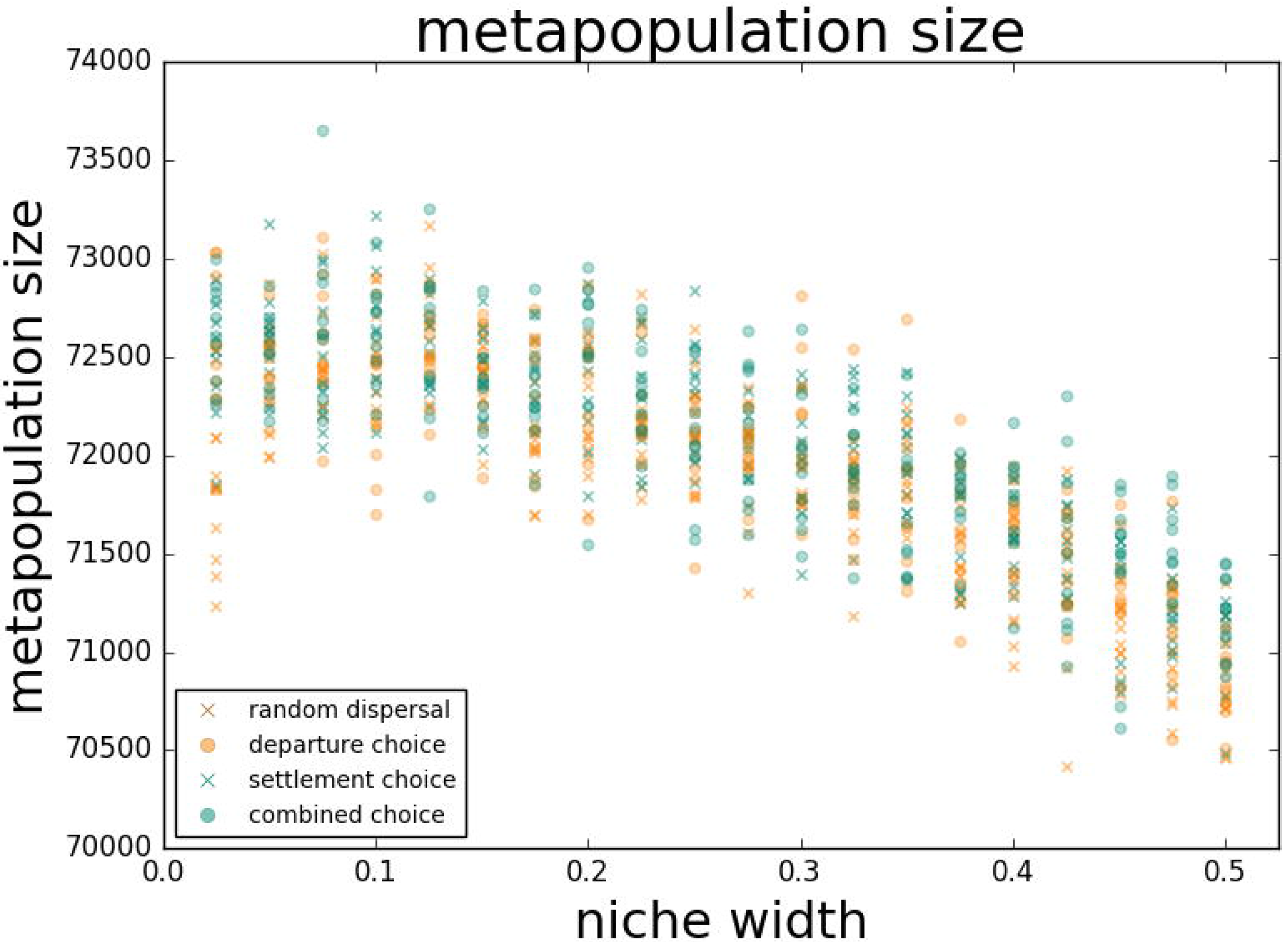
Metapopulation size for scenarios with fixed niche width. Scenarios of random (×) and informed (○) departure combined with random (orange) or informed (green) settlement.

**Figure 7:**
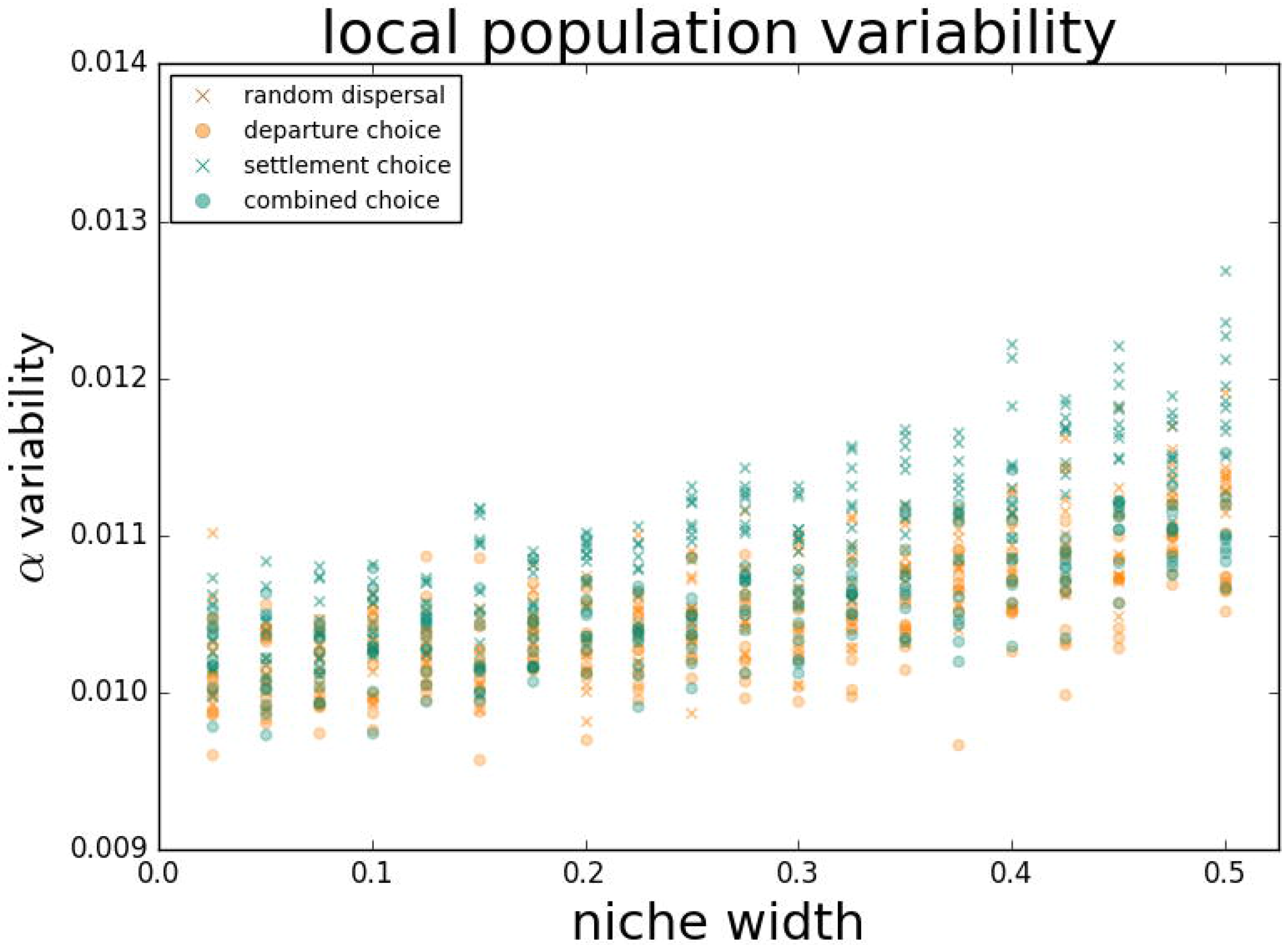
Local population variability for scenarios with fixed niche width. Scenarios of random (×) and informed (○) departure combined with random (orange) or informed (green) settlement.

### Imperfect habitat choice

For imperfect habitat choice, we found patterns intermediate to those of random and perfectly informed scenarios (see *supplementary material* 2), except in metapopulations of generalists that disperse most when having imperfect information.

## Discussion

Altering dispersal from a random process to one with habitat choice enables a reconciliation between ecological specialisation and dispersal evolution, as predicted. By allowing the evolution of ecological specialisation, habitat choice furthermore causes local populations to fluctuate less and metapopulation to reach higher sizes. Metapopulation-level performance is thus enhanced by habitat choice.

In accordance with previous studies, random dispersal leads to a trade-off between dispersal and ecological specialisation in the absence of habitat choice. Habitat choice changes this relationship to favour specialists even with high levels of dispersal and favour dispersal even in specialists. Our result confirms predictions from early theory [25, 45], as well as conclusions from more dedicated models with habitat choice either at departure [35, 36] or at settlement [32–34]. We take this one step further by contrasting departure and settlement decisions and demonstrating that choice mechanisms at settlement favour specialized strategies for even higher levels of dispersal than choice mechanisms at departure. A probable explanation for this stronger effect of settlement decision is that habitat choice at settlement integrates information from all potential settlement locations, rather than only from the natal location with informed departure. We show that a departure choice enables the evolution of even more specialized strategies when specialists are favoured. Dispersal from an already suitable habitat is costlier in more specialized strategies. Informed departure -i.e. informed philopatry-benefits specialists most by retaining individuals in their suitable habitat and enabling individuals in unsuitable habitat to disperse. Similar patterns were found in studies on habitat autocorrelation (discussed in [9]). When heterogeneity in habitat is experienced as coarse-grained relative to the scale of dispersal, the imposed landscape structure is more likely to match individuals to a suitable habitat, but when it is fine-grained, mismatches tend to prevail. Habitat matching by the spatial structure, instead of habitat choice behaviour, can limit maladaptive gene flow and select for increased specialism [9]. No matter how habitat matching is attained, we suspect that restricting movement of phenotypes to similar types of habitat always has the potential to favour specialism.

Habitat choice enables the evolution of dispersive specialists and altered metapopulation dynamics. Our model shows that metapopulation size is directly determined by the level of specialisation and only indirectly by information use during departure or settlement. This explains why the relation between metapopulation sizes and dispersal is inverse to that between evolved niche width and dispersal in scenarios of fixed dispersal. When generalism is favoured -i.e. under high levels of random settlement- the optimal strategy results in smaller metapopulation sizes despite the unaltered resource availability in our model. Individual interests do not necessarily line up with those of the metapopulation [46]. In addition, we show that local population sizes fluctuate more in metapopulations with more generalists, which also have smaller metapopulation sizes. Small metapopulations and high local population variability (α), which lead to more frequent occurrences of small local populations, increase the risk of extinction via demographic and genetic stochasticity [47, 48]. In favouring specialisation, habitat choice indirectly affects ecological dynamics, alters eco-evolutionary feedbacks and, as a consequence, can affect metapopulation persistence.

By demonstrating the impact of habitat choice on eco-evolutionary dynamics, our results question the validity of the numerous models assuming random dispersal [34, 49]. Dispersal without control during any phase likely applies to very few real systems such as bacteria and wind-dispersed plants. Many organisms are capable of more than just non-random dispersal, and move in a selective way, illustrated by obvious examples of habitat choice in grasshoppers [27, 28], cue-based preferences during movement [50] and habitat choice [24,51,52]. Selective movement is even found in organisms for which it seems less obvious, such as zoochorous plants that disperse their seeds to suitable habitat via animals [53] or plankton that drift on currents but are able to select where to settle [54, 55]. With the effect of habitat choice on ecological dynamics in mind, we recommend considering habitat choice when predicting the dynamics of spatially structured populations [23,34,42,49] -e.g. species distribution models, meta-community models, viability models. Our results show that in systems with habitat choice, assuming random dispersal may overlook habitat-choice effects on ecological specialisation, metapopulation size and stability.

Information use during habitat choice is assumed to be perfect in our model. This is a simplified representation of reality since very accurate information is costly (e.g. prospecting multiple locations or developing elaborate sensory organs; discussed in [42,55,56]) and, as a consequence, accuracy of information can itself be a subject of selection. Organisms should rarely be able to acquire high-precision information because of its high cost while a minimum of information is expected to be adaptive [57]. We present results for a scenario of perfect information use, but even under imperfect habitat choice our general evolutionary and ecological results stand (see *supplementary material 2*).

Different habitat-based choice behaviours that result in non-random dispersal may or may not differ in their eco-evolutionary consequences [31, 42]. Here, we show how the consequences of habitat choice at departure differ from those of habitat choice at settlement. Habitat choice may also vary in several other aspects, such as choice mechanisms and the reliability of information used [42, 58]. We expect such different variations on habitat choice to have additional influences on evolutionary and ecological processes of varying magnitude. Hence, disregarding these nuances might conceal some crucial insights or restrict us from generating detailed predictions. Unfortunately, we often lack information on the specifics of habitat choice in real-life populations. Our focus going forward should be on revealing the extent to which habitat choice varies along all these axes in nature, but also to evaluate their relative importance.

In summary, we demonstrate the profound effect of habitat choice on eco-evolutionary dynamics of metapopulations, including the dynamics of dispersal and ecological specialisation. Moreover, habitat choice at settlement impacts the model’s outcome more than choice at departure. Based on the difference between random dispersal and habitat choice, we encourage studies of real-world metapopulations to carefully consider habitat choice during the different phases of dispersal. Our results elucidate the impact of the often erroneous assumption of random dispersal and may improve the accuracy of future predictions of spatially structured populations.

## Acknowledgements

We would like to thank Pim Edelaar, François Massol and Estelle Laurent for engaging in insightful discussions. The project received financial support from the Research Foundation - Flanders (FWO) through research grant G.018017.N (FM, MVG & DB) and the research network EVENET (DB, SJ).

---

Table 1: fixed model parameters

